# Muntjac chromosome evolution and architecture

**DOI:** 10.1101/772343

**Authors:** Austin B. Mudd, Jessen V. Bredeson, Rachel Baum, Dirk Hockemeyer, Daniel S. Rokhsar

## Abstract

Despite their recent divergence, muntjac deer show striking karyotype differences. Here we describe new chromosome-scale genome assemblies for the Chinese and Indian muntjacs, *Muntiacus reevesi* (2n=46) and *Muntiacus muntjak* (2n=6/7), and analyze their evolution and architecture. We identified six fusion events shared by both species relative to the cervid ancestor and therefore present in the muntjac common ancestor, six fusion events unique to the *M. reevesi* lineage, and twenty-six fusion events unique to the *M. muntjak* lineage. One of these *M. muntjak* fusions reverses an earlier fission in the cervid lineage. Although comparative Hi-C analysis revealed differences in long-range genome contacts and A/B compartment structures, we discovered widespread conservation of local chromatin contacts between the muntjacs, even near the fusion sites. A small number of genes involved in chromosome maintenance show evidence for rapid evolution, possibly associated with the dramatic changes in karyotype. Analysis of muntjac genomes reveals new insights into this unique case of rapid karyotype evolution and the resulting biological variation.

## Background

Rapid karyotype evolution, or chromosomal tachytely [1], has been found in various species, such as rodents [2], bears [3], and gibbons [4], and as a byproduct of chromosomal instability in cancer [5]. Perhaps the most spectacular example of rapid karyotype evolution is found in muntjacs, a genus of small deer with karyotypes ranging from 2n=46 for *Muntiacus reevesi* to 2n=6/7 for female/male *Muntiacus muntjak*, respectively, with *M. muntjak* having the smallest chromosome number of any mammal [6]. Cytogenetic analysis showed that muntjac karyotype diversity arose primarily through centromere-telomere (head-tail) tandem fusions and, to a lesser extent, centromere-centromere (head-head) tandem fusions (*i.e.*, Robertsonian translocations [7]) [8, 9]. Importantly, independent fusions occurred in each lineage after divergence from their common ancestor, such that the 2n=46 *M. reevesi* karyotype does not represent an intermediate stage between the ancestral 2n=70 cervid karyotype and the highly reduced *M. muntjak* karyotype [10, 11].

Understanding the variation of genomic architectures in muntjacs has the potential to reveal new insights into chromosome evolution [12]. We therefore set out to explore karyotype changes in muntjacs by determining the number, distribution, and timing of shared and lineage-specific fusion events. To this end, we described the first chromosome-scale assemblies of *M. muntjak* and *M. reevesi* with contiguity metrics that surpass those of earlier draft assemblies [13, 14]. To infer the series of karyotype changes in muntjac, we leveraged existing assemblies of *Bos taurus* (cow) [15], *Cervus elaphus* (red deer) [16], and *Rangifer tarandus* (reindeer) [17]. In total, we characterized thirty-eight muntjac fusion events, six of which are shared by *M. muntjak* and *M. reevesi*. The rate of twenty-six unique fusion events in the *M. muntjak* lineage over 4.9 million years represents more than an order of magnitude increase relative to the mammalian average. Although the molecular mechanism driving these karyotype changes is unknown, we found that one fusion event in the *M. muntjak* lineage reversed a chromosome fission that occurred earlier in the cervid lineage; in another case, we found that a pair of ancestral cervid chromosomes likely fused independently in the *M. muntjac* and *M. reevesi* lineages. These findings suggest that some chromosomes may be more prone to karyotype changes than others and that care should be taken in applying the parsimony principle due to the possibility of convergent change.

We also took advantage of the extensive collinearity of the muntjac genomes to study changes in three-dimensional genome architecture that accompany chromosome fusions. Our findings suggest that while karyotype changes disrupt long-range three-dimensional genome structure, including A/B compartments, there are few changes at the local level. These analyses explore features of chromosome structure derived from the unique evolutionary history of these two karyotypically divergent species.

## Results and discussion

### Assembly and annotation

To investigate the tempo and mode of muntjac chromosome evolution, we generated high-quality, chromosome-scale genome assemblies for *M. muntjak* and *M. reevesi* (Table S1) using a combination of linked-reads (10X Genomics Chromium Genome) and chromatin conformation capture (Dovetail Genomics Hi-C; Table S2, Methods). The resulting assemblies each contain 2.5 Gb of contig sequence with contig N50 lengths over 200 kb. In both assemblies, over 92% of contig sequence is anchored to chromosomes. Compared with the publicly available assemblies [14], the assemblies described here represent a hundredfold improvement in scaffold N50 length and severalfold improvement in contig N50 length. As typical for short-read assemblies, our muntjac assemblies are largely complete with respect to genic sequences (see below) but are likely to underrepresent repetitive sequences such as pericentromeric heterochromatin and repetitive subtelomeric regions.

The assembled chromosome numbers recapitulate the karyotypes reported in the literature (2n=6 for female *M. muntjak* [18] and 2n=46 for *M. reevesi* [19]). *M. reevesi* chromosomes were validated against previously published chromosome painting data [20]. For *M. muntjak*, we aligned 377 previously sequenced bacterial artificial chromosomes (BACs) [21–23] and, based on corresponding fluorescent *in situ* hybridization (FISH) location data, found that 360 (95%) of BACs align to the expected chromosomes. Of the 17 BACs that align to a different chromosome than expected by FISH, 16 are well-aligned to our assembly in regions of conserved colinearity among cow, red deer, and muntjac chromosomes, which suggests that the FISH-based chromosome assignments of these BACs are likely incorrect. Only one of these 17 BACs aligns to two of our assembled *M. muntjak* chromosomes, indicating a possible local misassembly or BAC construction error.

For each muntjac genome, we annotated ∼26,000 protein-coding genes based on homology with *B. taurus* [15], *Ovis aries* (sheep) [24], and *Homo sapiens* (human) [25]. Over 98% of annotated genes are functionally annotated by InterProScan [26]. From these annotations, we identified 19,649 one-to-one gene orthologs between the two muntjac species as well as 7,953 one-to-one gene orthologs present in the two muntjacs, *B. taurus* [15], *C. elaphus* [16], and *R. tarandus* [17]. These ortholog sets were used in the evolutionary and phylogenomic analyses below (Figure 1A and 1C, Table S3, Methods). Gene set comparisons (Figure S1) show that the muntjac annotations include several thousand more conserved cervid genes than are found in the *C. elaphus* and *R. tarandus* annotations and demonstrate comparable completeness to *B. taurus*, supporting the completeness and accuracy of the muntjac assemblies in genic regions.

**Figure 1.**
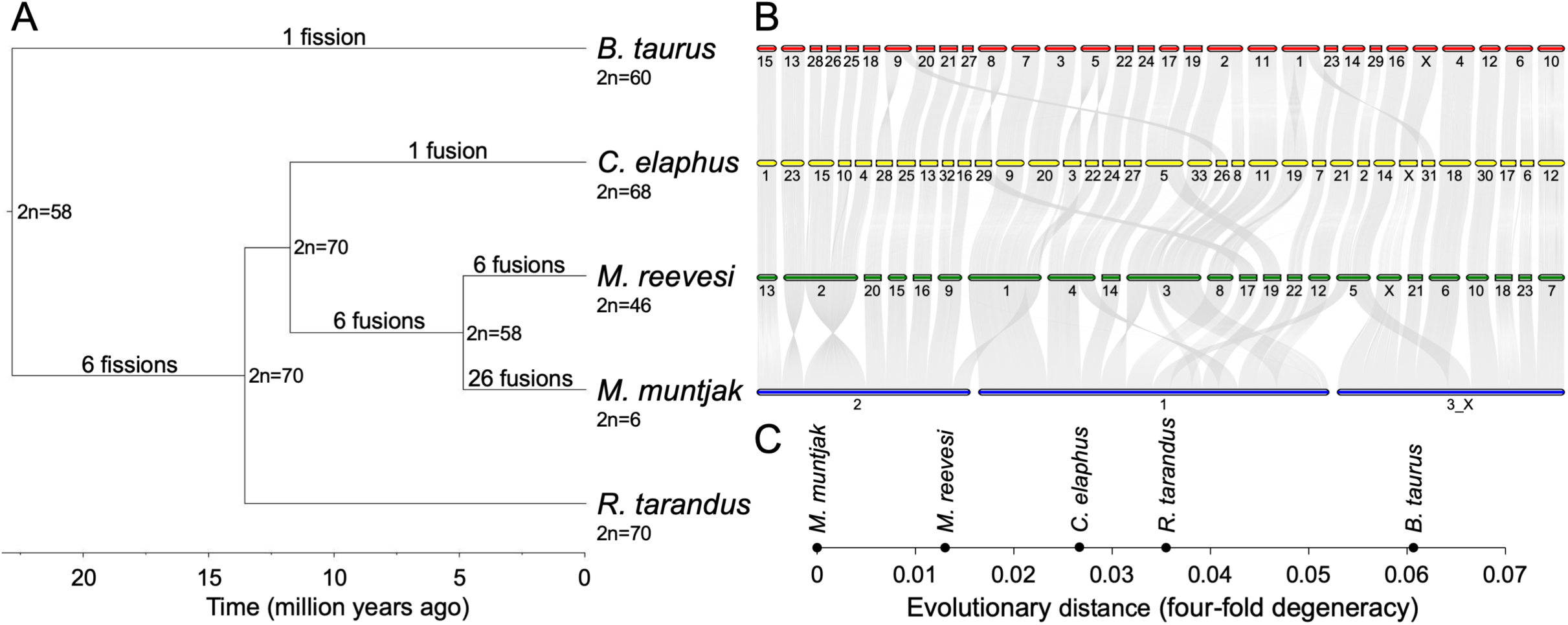
Evolutionary and phylogenomic analyses. [A] The phylogenetic tree of the five analyzed species, calculated from four-fold degenerate sites and divergence time confidence intervals, was visualized with FigTree (commit 901211e; https://github.com/rambaut/figtree). The tree denotes the ancestral karyotype at each node and the six branches with fission and fusion events relative to the ancestral karyotype. The lack of fissions or fusions on the *R. tarandus*-specific branch as well as the timings of the cervid-specific and *B. taurus*-specific fissions are derived from literature [20]. [B] Plot with jcvi.graphics.karyotype (v0.8.12; https://github.com/tanghaibao/jcvi) using runs of collinearity containing at least 25 kb of aligned sequence between *B. taurus*, *C. elaphus*, *M. reevesi*, and *M. muntjak*. *R. tarandus* was excluded, as it is not a chromosome-scale assembly. [C] Pairwise distances in substitutions per four-fold degenerate site extracted from the RAxML (v8.2.11) [66] phylogenetic tree using Newick utilities (v1.6) [63] are shown from reference genome *M. muntjak*.

### Comparative analysis

In order to study sequence and karyotype evolution, we aligned the two muntjac assemblies to each other and to *B. taurus* [15] as well as *B. taurus* to *C. elaphus* [16] and *R. tarandus* [17]. The pairwise alignment of the muntjac genomes contains 2.45 Gb of contig sequence, or over 97% of the assembled contig sequence lengths, with an average identity of 98.5% (excluding indels), reflecting the degree of sequence conservation between the two species and their recent divergence. In comparison, alignments of red deer, reindeer, and muntjacs to *B. taurus* contain 1.80 to 2.21 Gb of contig sequences with 92.7% to 93.2% average identity. Sequence alignments formed long runs of collinearity, and analysis of these alignments revealed the timing of fission and fusion events in each lineage (Figures 1A–B and S2A–D).

### Chromosome evolution

We assessed chromosome evolution in muntjacs using *B. taurus* (BTA) and *C. elaphus* (CEL) as outgroups. For convenience, we refer to chromosomal regions by their *B. taurus* (BTA) chromosome identifiers. We confirmed prior reports in literature [20] that:

1. In the last common ancestor of cow and deer, segments corresponding to the two cow chromosomes BTA26 and BTA28 were present as a single chromosome in the last common ancestor of cervids and *B. taurus*. This ancestral state, corresponding to BTA26_28, is retained in *C. elaphus* and the muntjacs.
2. Twelve chromosomes of the cervid ancestor arose by fission of chromosomes represented by six cow chromosomes (BTA1 => CEL19 and CEL31; BTA2 => CEL8 and CEL33; BTA5 => CEL3 and CEL22; BTA6 => CEL6 and CEL17; BTA8 => CEL16 and CEL29; and BTA9 => CEL26 and CEL28).
3. Although chromosomes homologous to BTA17 and BTA19 are fused in the *C. elaphus* lineage as CEL5, this fusion is unique to the *C. elaphus* lineage, and these cow chromosomes correspond to distinct ancestral cervid chromosomes.

In the muntjacs, we found six fusions shared by *M. muntjak* and *M. reevesi* (BTA7/BTA3, BTA5prox/BTA22, BTA2dist/BTA11, BTA18/BTA25/BTA26_28 (fusion of three ancestral chromosomes counted as two fusion events), and BTA27/BTA8dist; Figure S3). All six of these fusions shared by *M. muntjak* and *M. reevesi* were also confirmed in previous BAC-FISH analyses of *Muntiacus crinifrons*, *Muntiacus feae*, and *Muntiacus gongshanensis* [27, 28]. After the divergence of *M. muntjak* and *M. reevesi*, each lineage experienced additional fusions. In the *M. reevesi* lineage, there were six fusions (BTA7_3/BTA5dist, BTA18_25_26_28/BTA13, BTA2prox/BTA9dist/BTA2dist_11, BTA5prox_22/BTA24, and BTA29/BTA16). In the *M. muntjak* lineage, the three chromosomes arose via twenty-six lineage-specific fusions:

- *M. muntjak* chromosome 1: BTA7_3/BTA5prox_22/BTA17/BTA2prox/BTA1dist/BTA29/ BTA8prox/BTA9dist/BTA19/ BTA24/BTA23/BTA14/BTA2dist_11,
- *M. muntjak* chromosome 2: BTA15/BTA13/BTA18_25_26_28/BTA9prox/BTA20/BTA21/ BTA27_8dist/BTA5dist, and
- *M. muntjak* chromosome 3: BTAX/BTA1prox/BTA4/BTA16/BTA12/BTA6prox/BTA6dist/ BTA10.

We note that while both *M. muntjak* and *M. reevesi* karyotypes include chromosomes that arose by fusion of BTA13 and BTA18_25_26_28, these events appear to have occurred independently. Consistent with our analysis, published BAC FISH mapping of *M. reevesi* against *M. crinifrons*, *M. feae*, and *M. gongshanensis* found different locations of *B. taurus* chromosomes 13 and 18_25_26_28 in the muntjac species [27, 28]. This supports our finding that these are independent, lineage-specific fusion events.

In total, we found thirty-eight fusion events and no fissions separating the two muntjac species (Figure 1A). All twelve of the *M. reevesi* fusions identified by our comparative analysis are confirmed by BAC-FISH [20], and seventeen of the *M. muntjak* fusions are confirmed [29]. The additional fusions found in our analysis were not assayed by prior BAC-based studies. Our results are also consistent with the BAC-FISH findings of Chi et al. [9]. The rates of karyotype changes based on fission and fusion events in muntjacs are higher than the mammalian average of 0.4 changes per million years [30]. The *M. muntjak* lineage, with six fission events and thirty-two fusion events over the past 22.8 million years since the cervid ancestor, averaged 1.7 events per million years. In the 4.9 million years since the divergence from *M. reevesi*, this rate has increased to 5.3 fusion events per million years, an order of magnitude greater than the mammalian average. The *M. reevesi* lineage, on the other hand, averaged 0.8 events per million years over the past 22.8 million years, with an accelerated rate of 1.2 events per million years over the past 4.9 million years. Although the calculated nucleotide divergence and time between the two muntjac species (Figure 1A and 1C, Table S3) mirrors the evolutionary distance between humans and chimpanzees [31, 32], this number of fusion events since the muntjac last common ancestor far exceeds the rate in the chimpanzee and human lineages since their respective last common ancestor (*i.e.*, a single fusion on the human lineage [33]).

### Reversal of a cervid-specific fission in *M. muntjak*

While analyzing the fission and fusion events, we discovered a fusion in *M. muntjak* that reverses, to the resolution of our assembly, the cervid-specific fission of the ancestral chromosome corresponding to BTA6 (Figure S4). To estimate the probability of such a reversion occurring by chance given the high rate of fusion in *M. muntjak*, we simulated a simplified model for karyotype change with four rules: (1) only one fission is allowed per chromosome; (2) all fissions occur first, followed by all fusions; (3) for each fission, a chromosome is chosen at random; and (4) for each fusion, chromosomes and chromosomal orientations are chosen at random. From a starting karyotype of n=29, representing the last common ancestor of cervids and *B. taurus* [20], we simulated the model of fissions and fusions to one million iterations per fission-fusion combination (Figure S5). The *M. muntjak* lineage, with six fissions and thirty-two fusions, had a 4% probability of at least one fusion reversing a prior fission. In comparison, the *C. elaphus* lineage, with six fissions and one fusion, had only a 0.13% probability of reversal by chance, and the *M. reevesi* lineage, with six fissions and twelve fusions, had a 1.5% chance of reversal. Given the large number of fusions in muntjacs, the probability of a chance reversal of a previous fission is small; however, it is plausible that the reversal was aided by unmodeled effects of differential chromosome fusion probability arising, for example, by chromosomal proximity in the nucleus. This analysis points to the importance of having multiple outgroups (here both *B. taurus* and *C. elaphus*) in phylogenetic analyses of karyotypes.

### Changes in three-dimensional genome structure after karyotype change

Despite the extensive fusions documented above for *M. muntjak* and *M. reevesi*, the genomes are locally very similar (98.5% identity in aligned regions and fourfold synonymous substitution rate of 1.3%). Our Hi-C chromatin conformation capture data allows us to examine the impact of these rearrangements on local (*i.e.*, within a megabase along the genome) and longer (*i.e.*, 5 megabase) length scales as chromosomal segments become juxtaposed in novel ways. Focusing first on the *M. muntjak* and *M. reevesi* lineage-specific fusion sites (Tables S4-7), we note the maintenance of distinct Hi-C boundaries in several examples, such as the junction between *M. muntjak* chromosomes X and 3 at 133 Mb on chromosome 3_X. Other fusion sites, however, show no notable difference from the rest of the genome in *M. muntjak*. As expected, *M. reevesi* shows a clear distinction between intra- and inter-chromosomal contacts, including across fusion sites in *M. muntjak* (Figure 2). To quantify the chromatin changes at these fusion sites, we divided the genomes into 1 Mb bins and compared normalized inter-bin Hi-C contact between bins 5 Mb apart in the two species, using the *M. muntjak* assembly as the backbone for comparison (Figure S7). Confirming the initial visual analysis, we found that most bins containing a fusion site have fewer long-range chromatin contacts in *M. reevesi* (averaging 0.16 ± 0.09 normalized contacts per bin) compared with *M. muntjak* (averaging 0.62 ± 0.35 normalized contacts per bin), though we identified bins with few contacts in both species (Figure S7).

**Figure 2.**
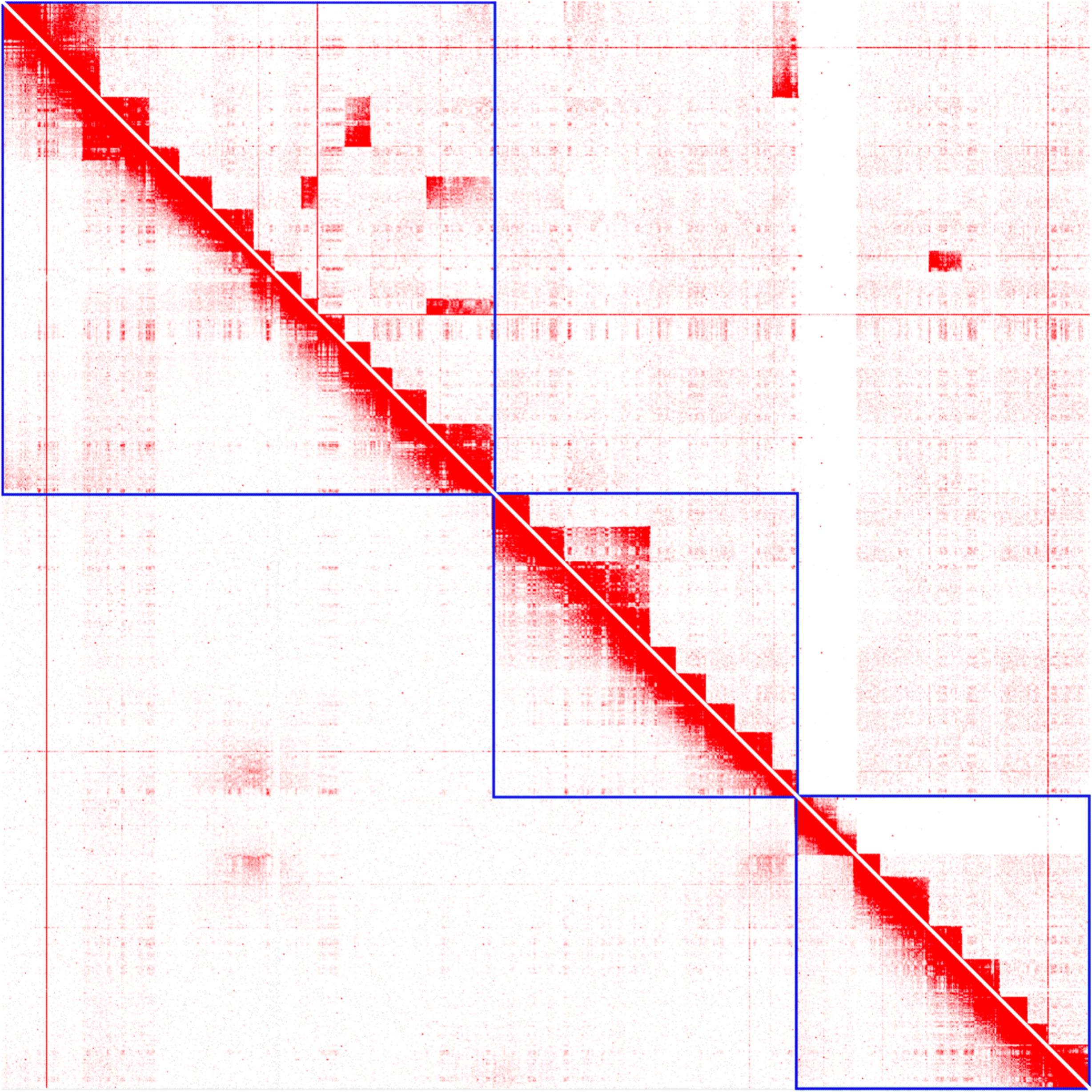
Chromosomal Hi-C contact maps. Visualization of the *M. muntjak* chromosomes’ Hi-C contact map (bottom left) and the *M. reevesi* chromosomes’ Hi-C contact map (top right) using the *M. muntjak* assembly as the reference in Juicebox (v1.9.0) [45]. The blue lines demarcate the boundaries of the three *M. muntjak* chromosomes.

In order to test whether differences are present at a more local level, we next compared normalized 1 Mb intra-bin Hi-C contacts between the two species, again using the *M. muntjak* assembly as the backbone for comparison. We found that most of the chromatin contacts are consistent between the two muntjacs, including all but three of the bins containing fusion sites (Figures 3A and S6). Several regions, however, show distinctive variation in chromatin contacts between the two species: the X chromosome and two regions on *M. muntjak* chromosome 1 (186–355 and 615–630 Mb). Since our sequenced *M. reevesi* sample is male [10] while the sequenced *M. muntjak* sample is female [34], we expect a difference in chromatin contacts on the X chromosome, a finding that is further supported by analysis of copy number across the genome using the 10X Genomics linked-read data (Figure 3B). From this copy number analysis, we also hypothesize that the two regions on *M. muntjak* chromosome 1 (186–355 and 615–630 Mb) are a haplotype-specific duplication and a haplotype-specific deletion, respectively, which would explain the difference in chromatin signal between the two muntjacs (Figure 3C–D). Although the inter-bin analysis identified long-range chromatin changes between sites 5 Mb apart, our quantitative comparison of 1 Mb intra-bin chromatin contacts found substantial chromatin conservation between the genome assemblies, including nearly all of the fusion sites. This conclusion is further supported by intra-bin analysis with 100 kb bins (Figure S8).

**Figure 3.**
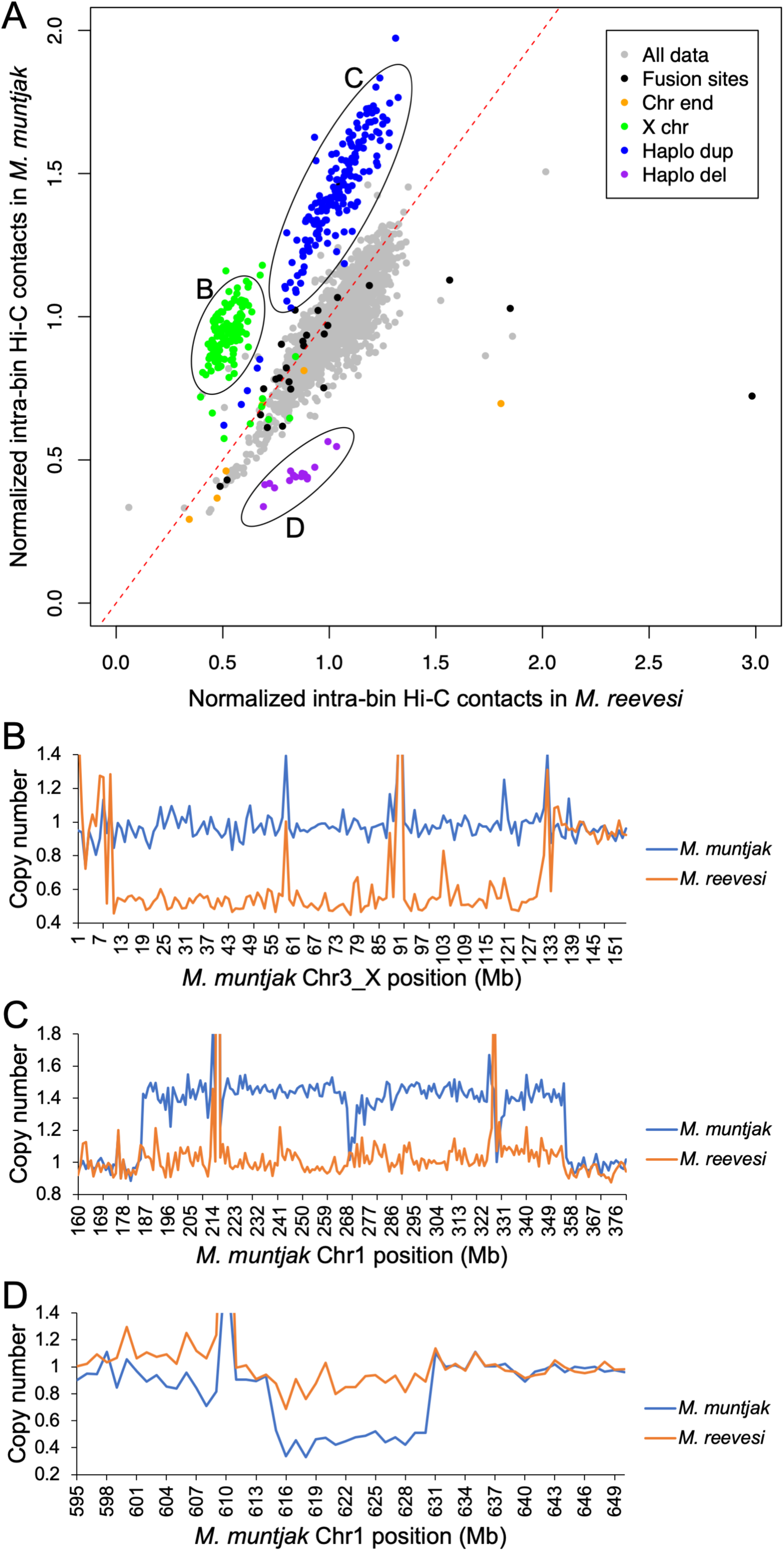
Evaluation of inter-chromosomal contacts. [A] 1 Mb intra-bin Hi-C contacts for *M. muntjak* (y axis) vs. *M. reevesi* (x axis) with the bins containing the *M. muntjak* lineage-specific fusion sites (Table S6), chromosome ends, the X chromosome, the potential *M. muntjak* haplotype-specific duplication, and the potential *M. muntjak* haplotype-specific deletion colored. The expected result of conserved Hi-C contacts is represented with a dashed red line. For fusion site ranges spanning two bins, the bin containing the majority of the fusion site range was deemed to be the fusion site bin. Copy number was calculated from normalized coverage of adapter-trimmed 10X Genomics linked-reads for three regions with variation in the chromatin contacts: [B] the X. chromosome, [C] the potential *M. muntjak* haplotype-specific duplication, and [D] the potential *M. muntjak* haplotype-specific deletion, with the copy number of *M. muntjak* in blue and *M. reevesi* in orange.

On a multi-megabase length scale, mammalian chromosomes can be subdivided into alternating A/B compartments based on intra-chromosome contacts; these compartments correspond to open and closed chromatin, respectively, and differ in gene density and GC content [35]. To test whether these compartments are conserved or disrupted by fusions, we computed the A/B chromatin compartment structures for *M. muntjak* and *M. reevesi* from the Hi-C data, again using the *M. muntjak* assembly as the backbone for comparison. We found that, in general, compartment boundaries are not well conserved between the muntjacs (Figure S9). Specifically, for A/B compartments larger than 3 Mb (*i.e*., containing more than three 1 Mb bins), only 17 compartments were completely conserved between the two species, out of 221 A/B compartments analyzed in *M. muntjak* and 161 in *M. reevesi*. We found that many of the compartments in *M. reevesi* are subdivided into multiple compartments in *M. muntjak*. Combining our analysis of A/B compartments and the chromatin contacts, we found that the extensive set of fusions in the *M. muntjac* lineage altered three-dimensional genome structure at the multi-megabase scale while still maintaining conservation at the local level. These large-scale chromatin changes accompanying karyotype change must have only limited effects on the underlying gene expression, since the two muntjac species can produce sterile hybrid offspring [36]. Similar uncoupling between genome topology and gene expression has been observed in *Drosophila melanogaster* [37].

### Genic evolution accompanying rapid karyotype change

Finally, we searched for genic differences between muntjac that may have accompanied rapid karyotype evolution. These could, for example, be mutations that led to dysfunctional chromosome maintenance and thus triggered the rapid occurrence of multiple fusions, such as by destabilization of telomeres. More subtly, these genic changes could have occurred as a response to chromosomal change; for example, the dramatic reduction in the number of telomeres following large-scale fusion could be permissive for mutations that make telomere maintenance less efficient. Our survey of gene and gene family differences between muntjacs were suggestive but ultimately inconclusive. In particular, we found evidence for positive selection of centromere-associated proteins CENPQ and CENPV and meiotic double strand break protein MEI4 as well as the expansion of the nucleosome-binding domain-containing HMG14 family in *M. muntjak*.

## Conclusions

We present here new chromosome-scale assemblies of two muntjac deer that differ dramatically in karyotype, despite only limited sequence change, after ∼4.9 million years of divergence. Analysis of these new assemblies revealed multiple changes in the underlying chromosome structure, including variation in the A/B compartments despite maintenance of local (*i.e.*, sub-megabase) three-dimensional genome contacts. One of the chromosome fusions reverses an earlier chromosome fission to the resolution of our assemblies, with the two events being separated by more than eight million years. Several chromosome maintenance associated proteins show accelerated evolution in *M. muntjak*, although functional studies will be required to determine any possible causal link to rapid karyotype change. Future studies will use these assemblies to resolve the nature of the fusion sites and to better understand the biological mechanisms related to chromosome fissions and fusions in muntjac.

## Methods

### DNA extraction and sequencing

High molecular weight DNA was extracted, as previously described [38], from fibroblast cell lines obtained from the University of Texas Southwestern Medical Center for *M. muntjak* (female) [34] and the University of Cambridge for *M. reevesi* (male) [10]. A 10X Genomics Chromium Genome library [39] was prepared for each species by the DNA Technologies and Expression Analysis Cores at the University of California Davis Genome Center and sequenced on the Illumina HiSeq X by Novogene Corporation. A Hi-C chromatin conformation capture library was also prepared for each species using the Dovetail Genomics Hi-C library preparation kit and sequenced on the Illumina HiSeq 4000 by the Vincent J. Coates Genomics Sequencing Laboratory at the University of California Berkeley.

### Shotgun assembly

10X Genomics linked-reads were assembled with Supernova (v2.0.1) [39]. Putative archaeal, bacterial, viral, and vector contamination was identified and removed by querying the assemblies using BLAST+ (v2.6.0) [40] against the respective RefSeq and UniVec databases, removing sequences with at least 95% identity, E-value less than 1E-10, and hits aligning to more than half the scaffold size or 200 bases, using custom script general_decon.sh (v1.0). Putative mitochondrial sequence was also identified and removed by querying the assemblies using BLAST+ (v2.6.0) [40] against their respective mitochondrial assemblies (NCBI NC_004563.1 [41] and NC_004069.1 [42]), removing sequence with at least 99% identity and E-value less than 1E-10, using custom script mt_decon.sh (v1.0). 71 scaffolds totaling 836 kb were removed from the *M. muntjak* assembly, and 36 scaffolds totaling 9 kb were removed from the *M. reevesi* assembly.

### Chromosomal assembly

Hi-C reads were aligned to each assembly with Juicer (v1.5.4-71-gd3ee11b) [43]. A preliminary round of Hi-C-based scaffolding was performed with 3D-DNA (commit 745779b) [44], and residual redundancy due to split haplotypes was manually filtered through visualization of the Hi-C contact map in Juicebox (v1.9.0) [45], removing the smaller of any pair of duplicate scaffolds. This process removed 1.04 Gb of sequence from the *M. muntjak* assembly and 25 Mb of sequence from the *M. reevesi* assembly. The remaining scaffolds were organized into chromosomes by realigning the Hi-C reads to the deduplicated assembly with Juicer (v1.5.4-71-gd3ee11b) [43], ordering and orienting scaffolds into chromosomes with 3D-DNA (commit 745779b) [44], and then manually correcting using Juicebox (v1.9.0) [45]. After correction, gaps in the assembly were filled with adapter-trimmed 10X Genomics data using custom script trim_10X.py (v1.0) and Platanus (v1.2.1) [46].

### Final assembly release and validation

Scaffolds smaller than 1 kb in the gap-filled assembly were removed with seqtk seq (v1.3-r106; https://github.com/lh3/seqtk), and chromosomes and scaffolds were numbered in order of size using SeqKit (v0.7.2-dev) [47]. X chromosomes were later renamed based on alignment with *B. taurus* [15]. Chromosomes in both species were oriented arbitrarily. For *M. reevesi*, the chromosome numbering in the assembly may differ from prior BAC-based studies. As *B. taurus* chromosome numbering is universally recognized, the extensive genomic collinearity of cervids, including both muntjacs, with cow provides a standard method of referencing homologous segments.

To validate the *M. muntjak* assembly, sequenced BACs [21–23] were aligned to it with BWA (v0.7.17-r1188) [48], and primary alignments were checked against the corresponding FISH locations, excluding unaligned BACs or those aligned to unplaced scaffolds.

### Annotation and homology analysis

Repetitive elements were identified and classified with RepeatModeler (v1.0.11) [49] and combined for each species with ancestral Cetartiodactyla repeats from RepBase (downloaded Nov 8, 2018) [50]. The assemblies were then soft masked with RepeatMasker (v4.0.7) [51]. The assemblies were annotated using Gene Model Mapper (v1.5.3) [52] and BLAST+ (v2.6.0) [40] with the following assemblies and annotations from Ensembl release 94 [53] as input evidence: *B. taurus* (Sep 2011 genebuild of GCA_000003055.3) [15], *H. sapiens* (Jul 2018 genebuild of GCA_000001405.27) [25], and *O. aries* (May 2015 genebuild of GCA_000298735.1) [24]. Coding nucleotide and peptide sequences were extracted using gff3ToGenePred and genePredToProt from the UCSC Genomics Institute (binaries downloaded 2019-03-05) [54] using custom script postGeMoMa.py (v1.0), and functional annotation was run with InterProScan (v5.34-73.0) [26].

Pairwise gene homology of the two muntjac annotations as well as total gene homology of the two muntjac, *B. taurus* (Ensembl release 94 Sep 2011 genebuild of GCA_000003055.3) [15, 53], *C. elaphus* (publication genebuild of GCA_002197005.1) [16], and *R. tarandus* (release date 2017-10-17 genebuild) [17] annotations were analyzed with OrthoVenn [55] using the default E-value of 1e-5 and inflation value of 1.5. One-to-one orthologous muntjac genes were extracted from the pairwise OrthoVenn output, and Yang-Nielsen synonymous and nonsynonymous substitution rates were calculated with the Ks calculation script (commit 78dda1e; https://github.com/tanghaibao/bio-pipeline/tree/master/synonymous_calculation) using ClustalW2 (v2.1) [56] and PAML (v4.7) [57]. Gene gain was identified from the full gene homology OrthoVenn output, requiring that the number of *M. muntjak* genes in an OrthoVenn cluster be greater than the number of genes found in any other analyzed species. Putative gene names of the results were extracted from the BLAST+ (v2.6.0) [40] best hit to the *H. sapiens* proteome from UniProt [58].

### Comparative analysis

The two muntjac assemblies were aligned to each other with cactus (v1590-ge4d0859) [59]. After removing any ambiguous sequence with seqtk randbase (v1.3-r106; https://github.com/lh3/seqtk), the muntjac assemblies, *C. elaphus* (GCA_002197005.1) [16], and *R. tarandus* (release date 2017-10-17 version) [17] were each also aligned pairwise against *B. taurus* (GCA_000003055.3) [15] with cactus (v1590-ge4d0859) [59]. Using custom script cactus_filter.py (v1.0), all pairwise output HAL alignment files were converted into PSL format with halLiftover (v200-gf7287c8) [60]. Using tools from the UCSC Genomics Institute (binaries downloaded 2019-03-05) [54] unless noted otherwise, the PSL files were filtered and converted with pslMap, axtChain, chainPreNet, chainCleaner (commit aacca59) [61], chainNet, netSyntenic, netToAxt, axtSort, and axtToMaf. Runs of collinearity were extracted from each pairwise MAF file by linking together local alignment blocks where the species 1 and species 2 locations, correspondingly, are in the same orientation and are neighboring in their respective genomes without intervening aligned sequence from elsewhere in the genomes. The pairwise MAF files from the alignments against *B. taurus* were also merged with ROAST/MULTIZ (v012109) [62], using the phylogenetic topology extracted with Newick utilities (v1.6) [63] from a consensus tree of the species from 10kTrees [64], and sorted with last (v912) [65].

### Phylogeny

From the one-to-one orthologous genes of all five species identified by OrthoVenn, codons with potential four-fold degeneracy were extracted from the *B. taurus* Ensembl release 94 Sep 2011 genebuild, excluding codons spanning introns, using custom script 4Dextract.py (v1.0). Using the ROAST-merged MAF file with *B. taurus* as reference, the corresponding codons were identified in the other four species, checking for corresponding amino acid conservation and excluding any codons that span two alignment blocks in the MAF file. The output fasta file containing four-fold degenerate bases was converted in phylip format with BeforePhylo (commit 0885849; https://github.com/qiyunzhu/BeforePhylo) and then analyzed with RAxML (v8.2.11) [66] using the GTR+Gamma model of substitution with outgroup *B. taurus*. As previously described [67], the divergence time confidence intervals from TimeTree [68] for all nodes except the outgroup *B. taurus* node were input into MEGA7 (v7.0.26) [69] using the Reltime method [70] and the GTR+Gamma model to create a time tree. To confirm the resulting times, the time calculated for the outgroup *B. taurus* node was verified in the literature [71].

### Chromatin conformation analysis

Hi-C reads from both species were aligned to the *M. muntjak* assembly with Juicer (v1.5.4-71-gd3ee11b) [43], and KR normalized intrachromosomal Hi-C contact matrices were extracted with Juicer Tools (v1.5.4-71-gd3ee11b) [43] at 1 Mb resolution. A sliding window-based localized principle component analysis (PCA) was used to call A/B compartment structure using custom script call-compartments.R (https://bitbucket.org/bredeson/artisanal). Localization of the PCA along the diagonal of the Pearson correlation matrix (forty 1 Mb windows with a step of twenty) mitigates confounding signal from large-scale intrachromosomal interarm contacts and amplifies compartment signal.

Hi-C contacts from the Juicer (v1.5.4-71-gd3ee11b) [43] merged_nodups.txt output file were split into 1 Mb and 100 kb bins using custom scripts HiCbins_1Mb.py and HiCbins_100kb.py, respectively. Intra-bin and inter-bin Hi-C contacts were extracted and normalized based on the average number of contacts per bin for each species.

### Copy number analysis

To explore the three regions with variation in chromatin contacts, adapter trimmed 10X Genomics data for each species was aligned to the *M. muntjak* assembly with BWA (v0.7.17-r1188) [48]. Alignment depth was extracted with SAMtools (v1.6) [72], and copy number was calculated from the average alignment depth per 1 Mb bin for each species.

## Abbreviations

B. Taurus: Bos Taurus
C. elaphus: Cervus elaphus
H. sapiens: Homo sapiens
M. crinifrons: Muntiacus crinifrons
M. feae: Muntiacus feae
M. gongshanensis: Muntiacus gongshanensis
M. muntjak: Muntiacus muntjak
M. reevesi: Muntiacus reevesi
O. aries: Ovis aries
R. tarandus: Rangifer tarandus
BAC: bacterial artificial chromosome
FISH: fluorescent in situ hybridization
prox: proximal
dist: distal
PCA: principle component analysis

## Declarations

### Ethics approval and consent to participate

Not applicable

### Consent for publication

Not applicable

### Availability of data and materials

The assemblies, annotations, and raw data for *M. muntjak* and *M. reevesi* were deposited at NCBI under BioProjects PRJNA542135 and PRJNA542137, respectively. Supporting files for the repeat and gene annotations are available at https://doi.org/10.6078/D1KT16. Unless otherwise stated, custom code used in this study is available at https://github.com/abmudd/Assembly.

### Authors’ contributions

ABM assembled and annotated the genomes, completed the bioinformatic analyses, and wrote the manuscript. JVB assisted with the bioinformatic analyses and script development. RB prepared the Hi-C libraries. DH coordinated the cell line acquisitions, extracted the DNA, and prepared the Hi-C libraries. DH and DSR provided scientific leadership of the project and wrote the manuscript. All authors reviewed the manuscript.

## Acknowledgements

We thank Jerry Shay and Woody Wright for providing the *M. muntjak* cell line; Malcolm Ferguson-Smith and Fengtang Yang for providing the *M. reevesi* cell line; Karen Lundy and the Functional Genomics Laboratory at the University of California Berkeley for running quality control on the extracted DNA; Dovetail Genomics for providing the Hi-C library preparation kit and running quality control on the Hi-C libraries; Shana McDevitt and the Vincent J. Coates Genomics Sequencing Laboratory at the University of California Berkeley for sequencing the Hi-C libraries; Jessica Lyons for coordinating the preparation and sequencing of the *M. muntjak* 10X Genomics library; Diana Burkart-Waco and the DNA Technologies and Expression Analysis Cores at the University of California Davis Genome Center for preparing the 10X Genomics libraries; Novogene Corporation for sequencing the 10X Genomics libraries; and Gary Karpen for providing comments on the manuscript.

---

**Figure S1.**
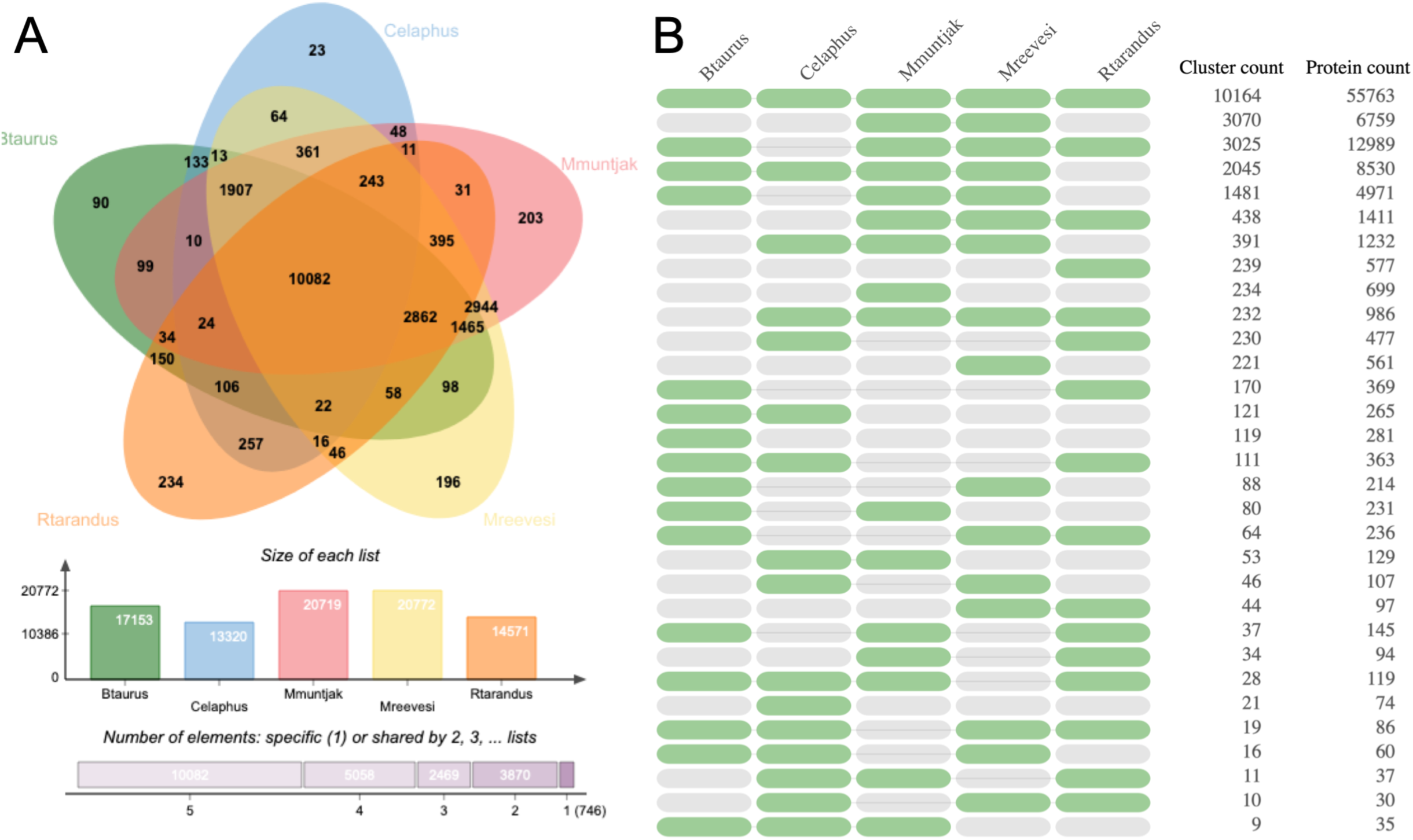
[A] Venn diagram of gene homology between the two muntjac annotations, *B. taurus* (Ensembl release 94 Sep 2011 genebuild of GCA_000003055.3) [15, 53], *C. elaphus* (publication genebuild of GCA_002197005.1) [16], and *R. tarandus* (release date 2017-10-17 genebuild) [17] annotations analyzed with OrthoVenn [55] and [B] the occurrence table of gene homology between these species reanalyzed with OrthoVenn2 [73] for visualization purposes. In the occurrence table, the green and grey represent the presence or absence, respectively, of that species in the OrthoVenn 2 clustering. The number of clusters and proteins are provided for all species combinations.

**Figure S2.**
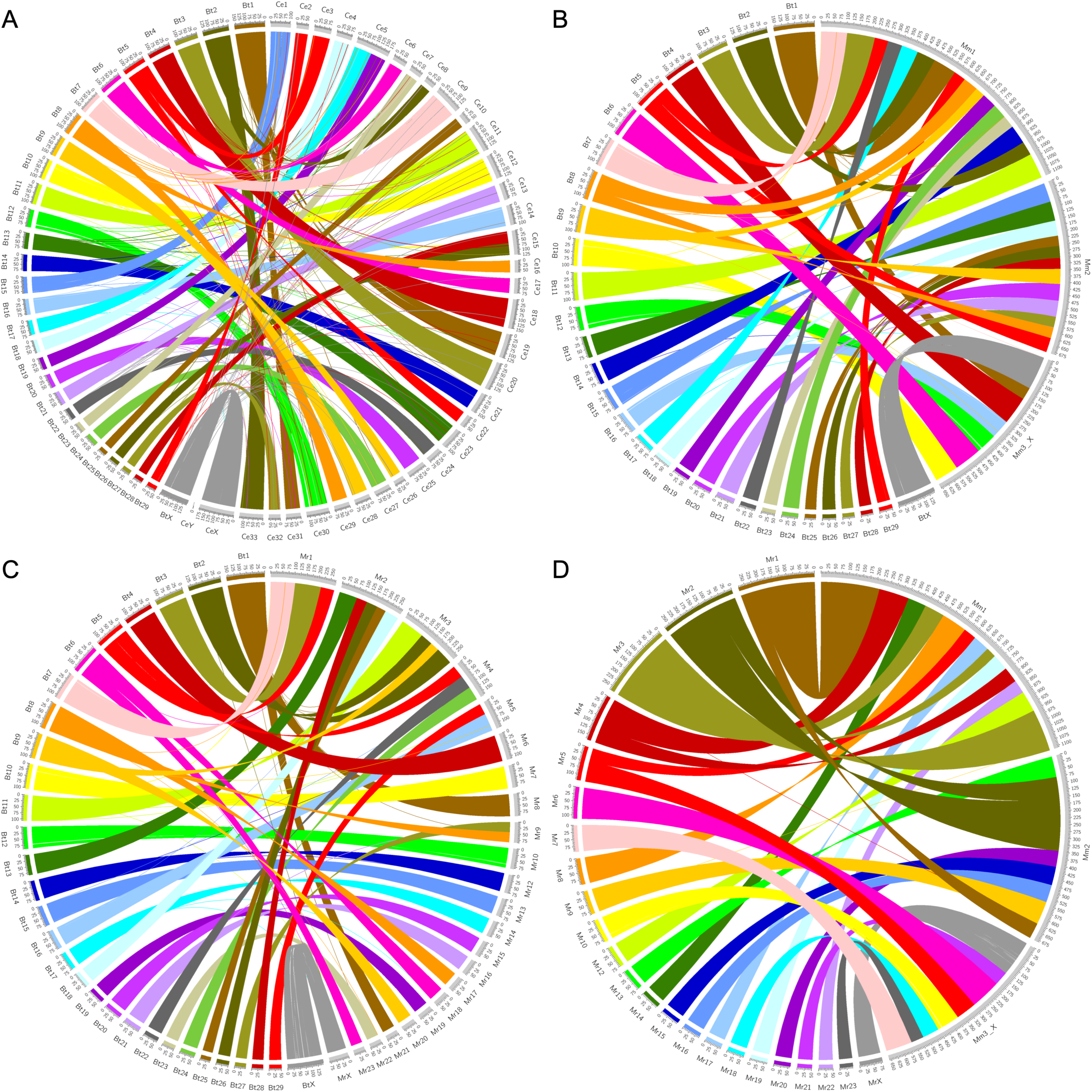
Circos (v0.69-6) [74] plots with runs of collinearity containing at least 25 kb of aligned sequence between [A] *B. taurus* (left, Bt) and *C. elaphus* (right, Ce), [B] *B. taurus* (left, Bt) and *M. muntjak* (right, Mm), [C] *B. taurus* (left, Bt) and *M. reevesi* (right, Mr), and [D] *M. reevesi* (left, Mr) and *M. muntjak* (right, Mm).

**Figure S3.**
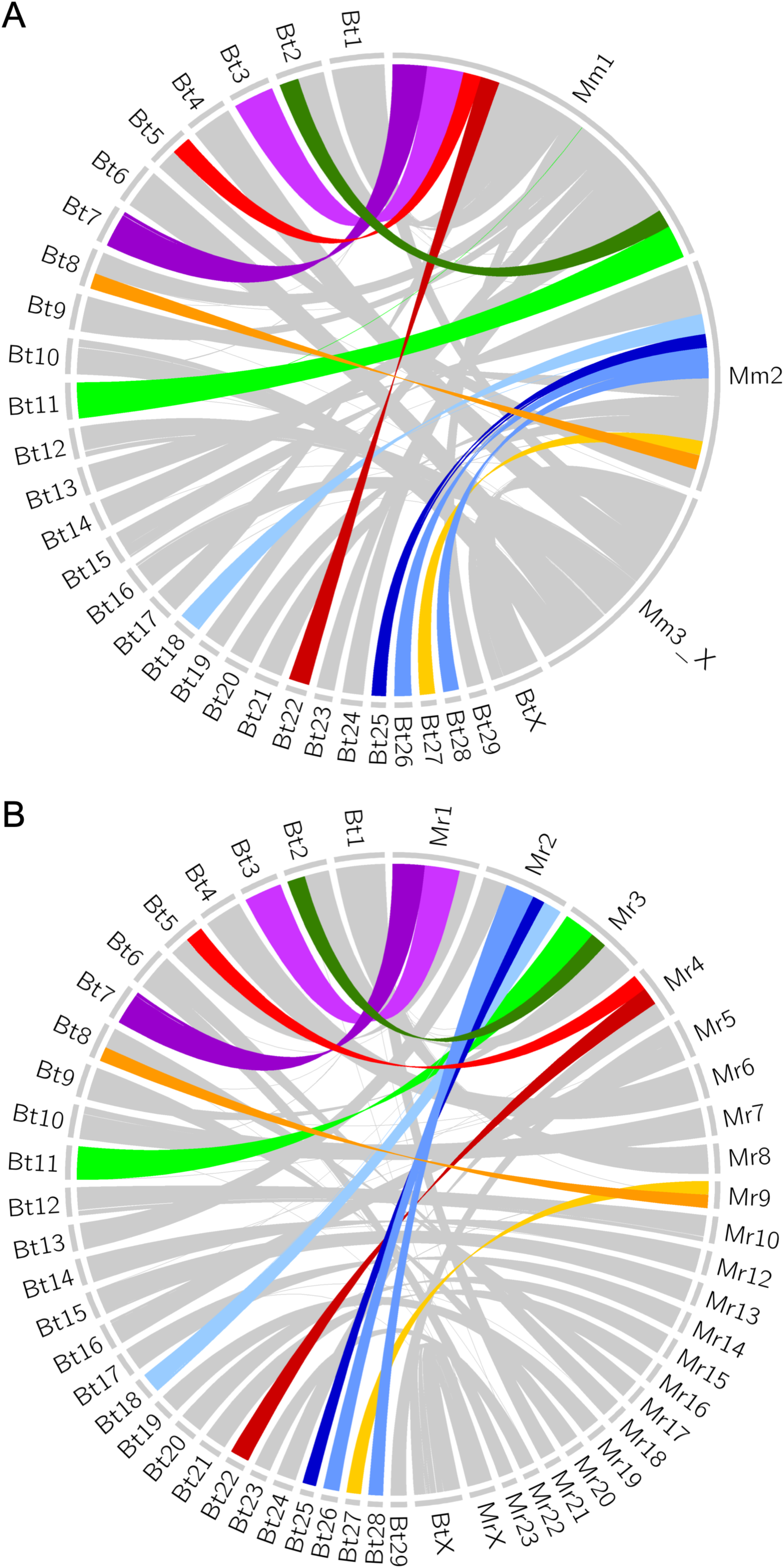
Circos (v0.69-6) [74] plots using runs of collinearity containing at least 25 kb of aligned sequence between [A] *B. taurus* (left, Bt) and *M. muntjak* (right, Mm) and [B] *B. Taurus* (left, Bt) and *M. reevesi* (right, Mr) specifying the six shared muntjac fusions: 7/3 (purple), 5prox/22 (red), 2dist/11 (green), 18/25/26_28 (blue), and 27/8dist (orange).

**Figure S4.**
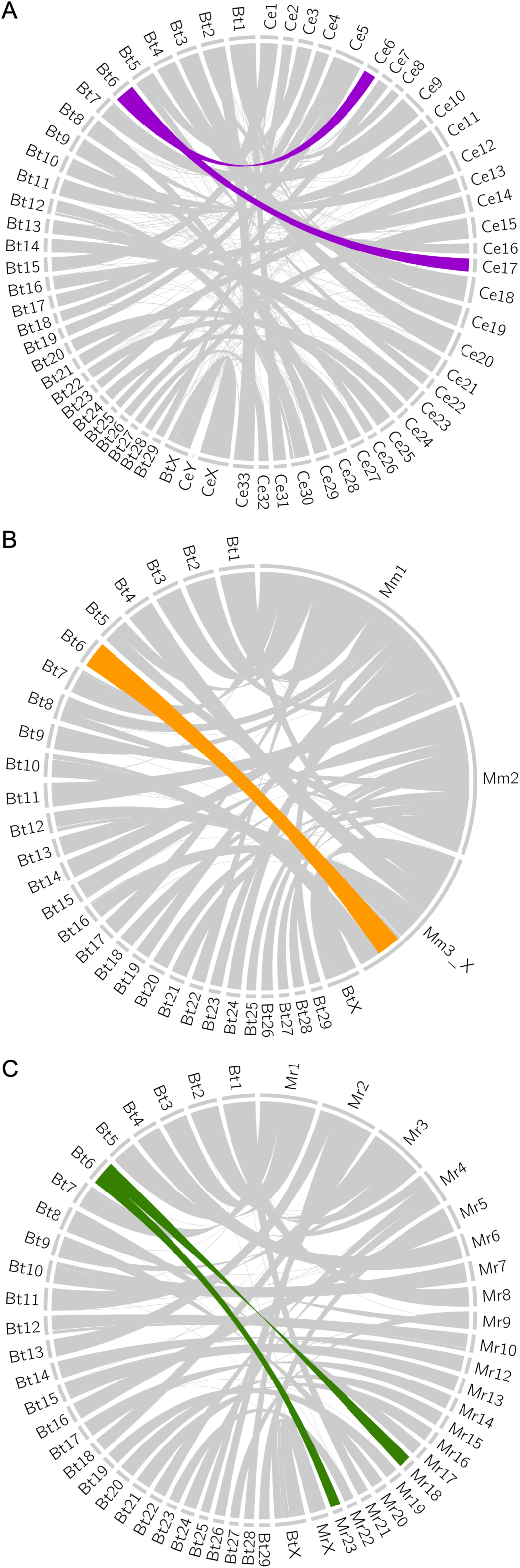
Circos (v0.69-6) [74] plots using runs of collinearity containing at least 25 kb of aligned sequence between [A] *B. taurus* (left, Bt) and *C. elaphus* (right, Ce) with the fission of *B. taurus* chromosome 6 in purple; [B] *B. taurus* (left, Bt) and *M. muntjak* (right, Mm) with the fission-fusion reversal of *B. taurus* chromosome 6 in orange; and [C] *B. taurus* (left, Bt) and *M. reevesi* (right, Mr) with the fission of *B. taurus* chromosome 6 in green.

**Figure S5.**
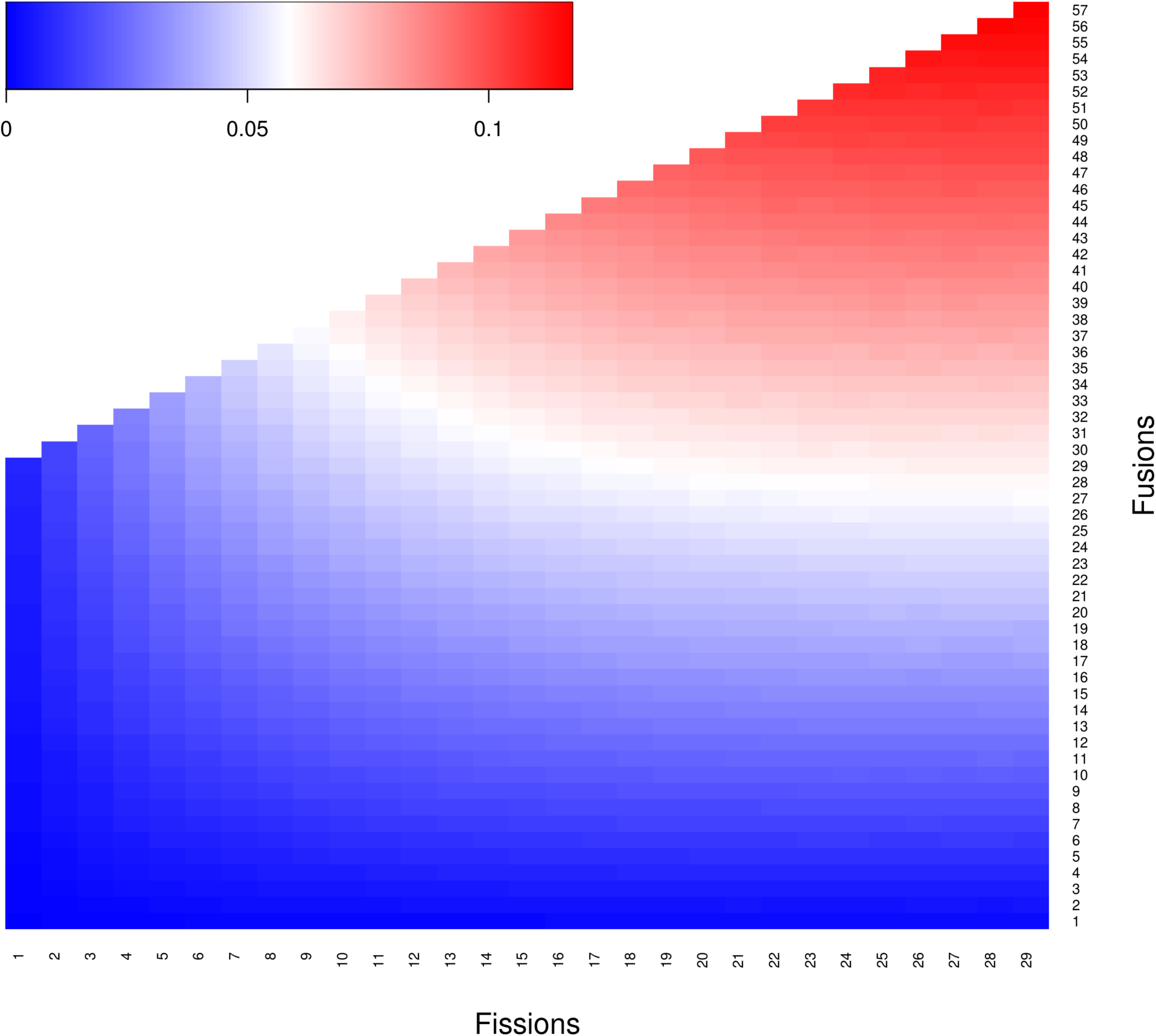
Heatmap of probabilities where at least one fusion reverses a prior fission modeled to one million iterations for each possible scenario from a starting karyotype of n=29 using custom script run_fission_fusion.sh (v1.0).

**Figure S6.**
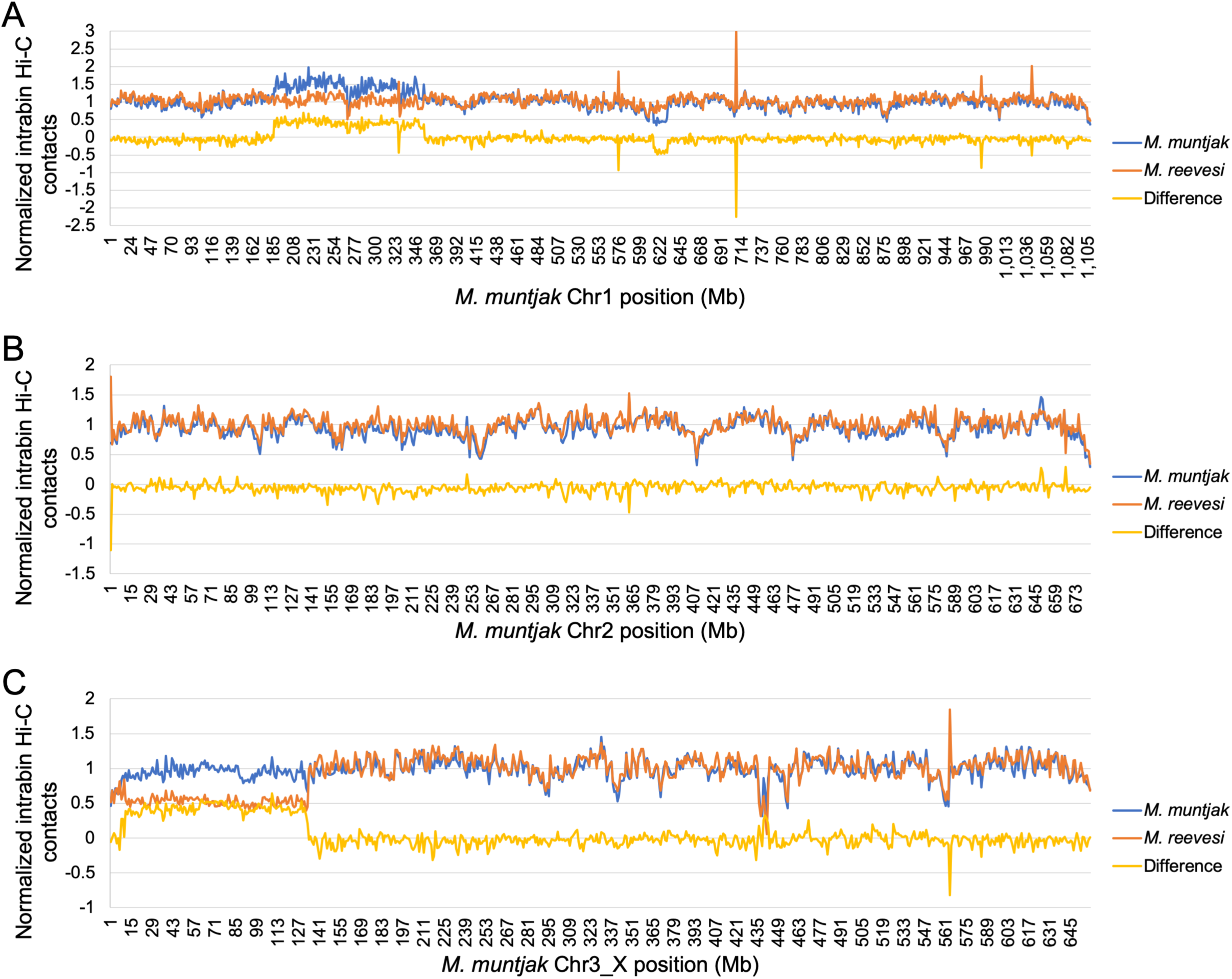
Using a bin size of 1 Mb and the *M. muntjak* assembly as the reference, normalized intra-bin Hi-C contacts for *M. muntjak* (blue) and *M. reevesi* (orange) at each position on [A] chromosome 1, [B] chromosome 2, and [C] chromosome 3_X. The difference of *M. muntjak* contacts minus *M. reevesi* contacts is displayed in yellow.

**Figure S7.**
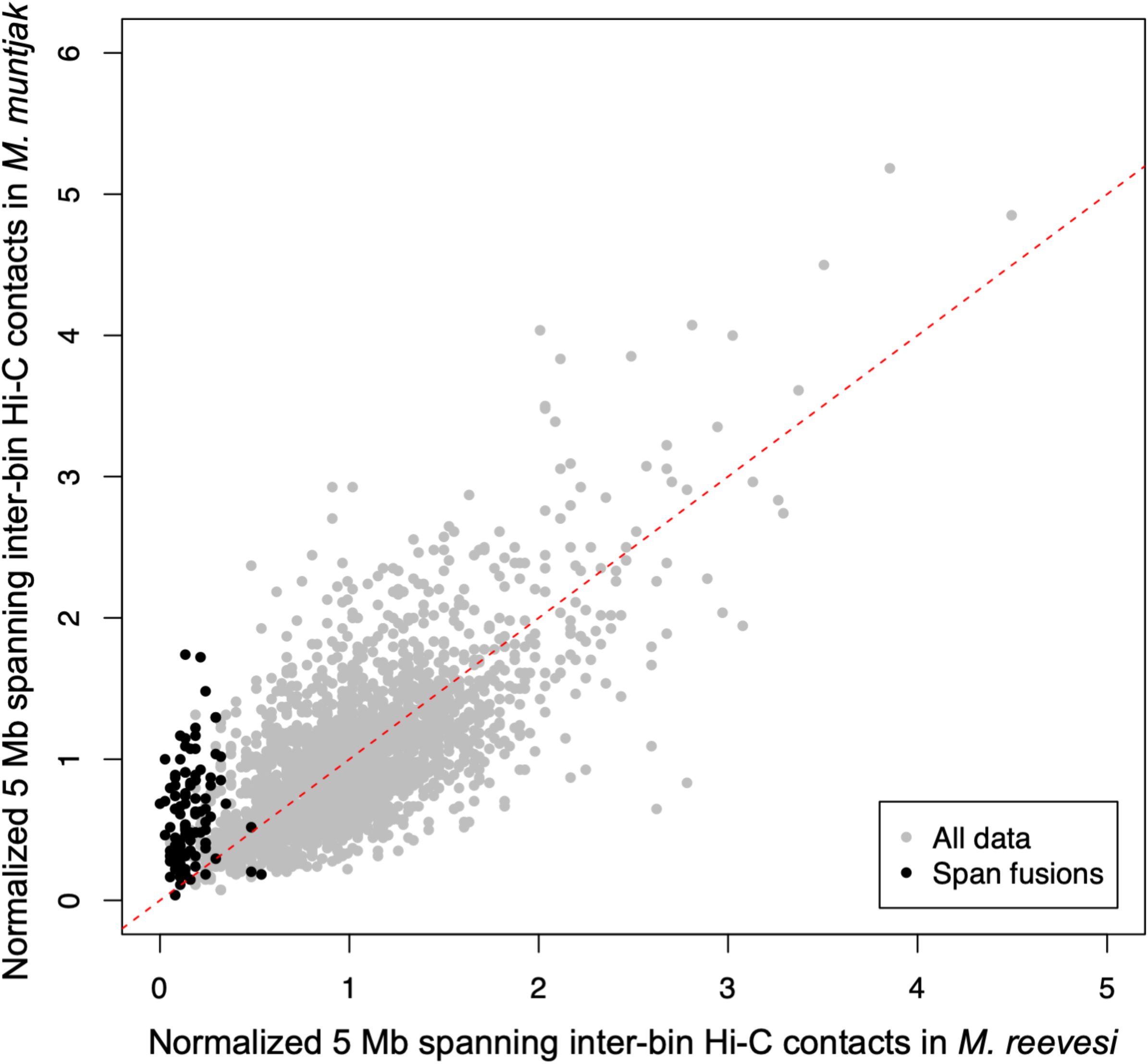
1 Mb inter-bin Hi-C contacts between bins 5 Mb apart for *M. muntjak* (y axis) vs. *M. reevesi* (x axis) with the inter-bin contacts that span across but do not include the *M. muntjak* lineage-specific fusion sites (Table S6) colored black. The expected result of conserved Hi-C contacts is represented with a dashed red line.

**Figure S8.**
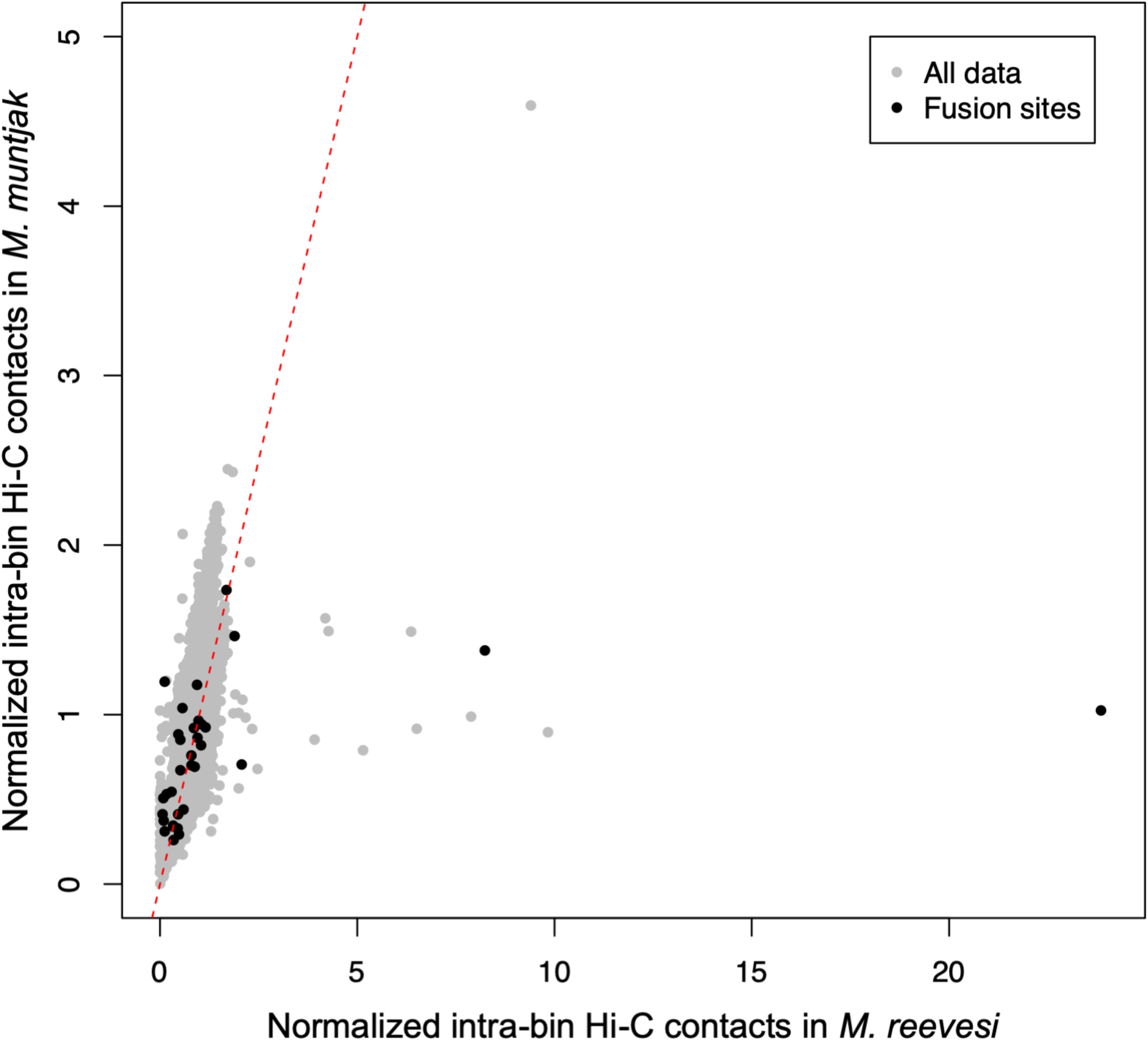
100 kb intra-bin Hi-C contacts for *M. muntjak* (y axis) vs. *M. reevesi* (x axis) with the bins containing the *M. muntjak* lineage-specific fusion sites (Table S6) colored black. The expected result of conserved Hi-C contacts is represented with a dashed red line. For fusion site ranges spanning two bins, the bin containing the majority of the fusion site range was deemed to be the fusion site bin. For fusion site ranges spanning three or more bins, the middle 100 kb bin(s) was deemed to be the fusion site bin(s).

**Figure S9.**
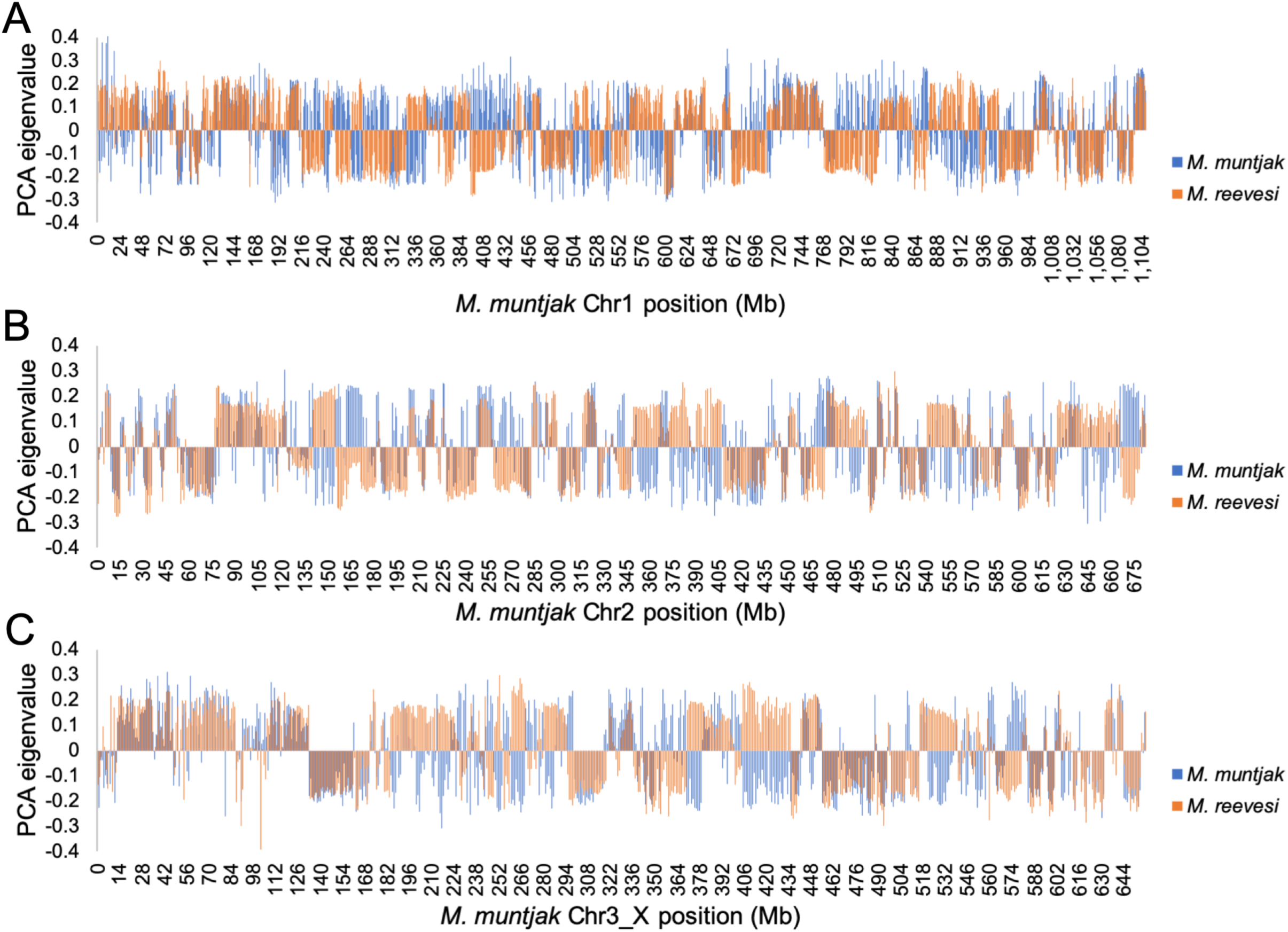
Using the *M. muntjak* assembly as reference, identification of A/B compartment boundaries for *M. muntjak* (blue) and *M. reevesi* (orange) based on PCA 1 eigenvalues with call-compartments.R.

**Table S1.** Summary of genome assembly. Statistics were calculated using assemblathon_stats.pl (commit d1f044b) [75] and GenomeTools (v1.5.9) [76].

| Genomic feature | *M. muntjak* | *M. reevesi* |
| --- | --- | --- |
| Total scaffold length, bp | 2,573,529,099 | 2,579,575,442 |
| Number of scaffolds | 25,651 | 29,705 |
| Scaffold N50 length, bp | 682,452,208 | 94,101,870 |
| Total contig length, bp | 2,518,738,577 | 2,514,747,046 |
| Number of contigs | 49,270 | 53,090 |
| Contig N50 length, bp | 215,534 | 225,142 |
| Contigs sequence in chromosomes, % | 95.06 | 92.93 |
| Contig GC content, % | 41.59 | 41.59 |
| Masked contig repeat sequence, % | 40.33 | 40.06 |
| Number of genes | 25,753 | 26,054 |
| Genes with functional annotation, % | 98.11 | 98.15 |
| Average number of exons per gene | 7.83 | 7.77 |
| Median gene size, aa | 328 | 326 |
| Median exon size, bp | 124 | 124 |
| Median intron size, bp | 921 | 911 |

**Table S2.**
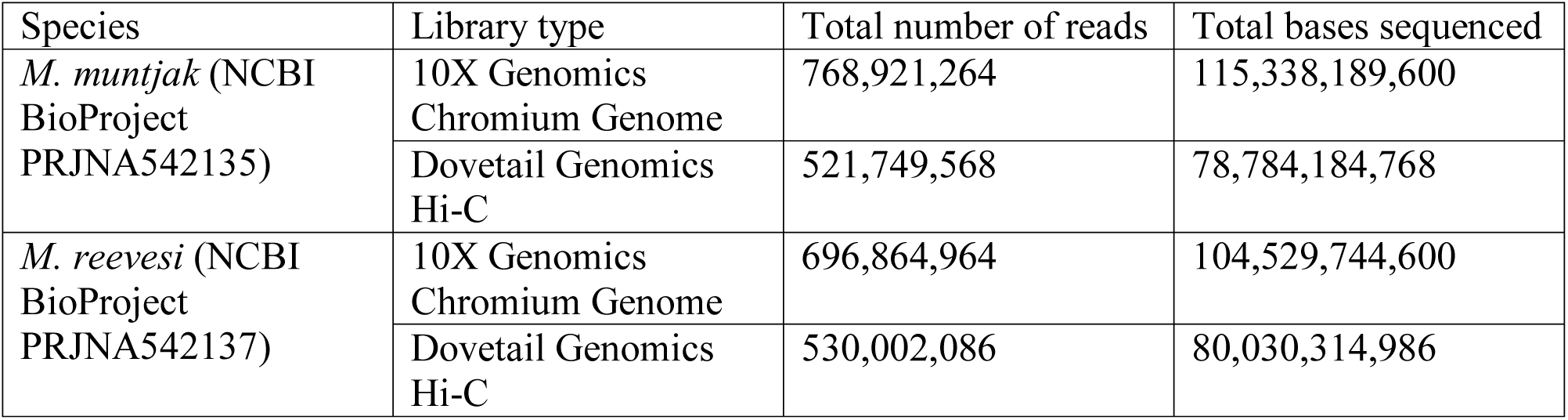
DNA sequencing generated for the genome assembly.

**Table S3.** Pairwise nucleotide divergence based on four-fold degeneracy between the examined species.

| Species | *B. taurus* | *C. elaphus* | *M. muntjak* | *M. reevesi* |
| --- | --- | --- | --- | --- |
| *C. elaphus* | 0.0549632 | – | – | – |
| *M. muntjak* | 0.0606332 | 0.0266608 | – | – |
| *M. reevesi* | 0.0598953 | 0.0259229 | 0.0130037 | – |
| *R. tarandus* | 0.0591649 | 0.0298125 | 0.0354825 | 0.0347446 |

**Table S4.** Locations of six cervid-specific fissions on the *B. taurus* genome assembly.

| *B. taurus* chromosome | Using runs of collinearity from *C. elaphus* | Using runs of collinearity from *M. muntjak* | Using runs of collinearity from *M. reevesi* |
| --- | --- | --- | --- |
| BTA1 | 58,941,477 – 58,978,602 | 57,645,593 – 57,778,258 | 57,645,547 – 57,746,127 |
| BTA2 | 93,282,776 – 93,424,724 | 79,668,309 – 79,719,766 | 79,668,935 – 79,719,765 |
| BTA5 | 70,623,938 – 70,699,763 | 57,880,818 – 58,822,584 | 57,880,818 – 60,196,482 |
| BTA6 | 63,301,740 – 63,370,450 | Fusion reversal | 68,435,554 – 68,455,313 |
| BTA8 | 64,071,291 – 64,114,095 | 67,266,180 – 67,499,444 | 67,369,805 – 67,497,566 |
| BTA9 | 63,670,677 – 64,013,115 | 64,824,832 – 65,087,945 | 64,824,832 – 65,087,945 |

**Table S5.** Locations of shared fusion events in the *M. muntjak* and *M. reevesi* genome assembly using runs of collinearity from *B. taurus*.

| *B. taurus* fused chromosomes | *M. muntjak* | *M. reevesi* |
| --- | --- | --- |
| 7/3 | Chr1: 103,142,901 – 103,201,151 | Chr1: 103,650,999 – 103,852,521 |
| 5prox/22 | Chr1: 267,859,762 – 267,926,350 | Chr4: 52,627,734 – 52,781,577 |
| 2dist/11 | Chr1: 1,006,153,244 – 1,006,638,886 | Chr3: 100,711,306 – 101,302,993 |
| 18/25 | Chr2: 216,351,804 – 216,390,051 | Chr2: 210,159,178 – 210,200,937 |
| 25/26 | Chr2: 256,956,082 – 257,281,016 | Chr2: 169,793,458 – 170,063,149 |
| 26_28 (*B. taurus* fission) | Chr2: 305,072,202 – 305,072,202 | Chr2: 121,918,267 – 122,082,397 |
| 27/8dist | Chr2: 580,274,743 – 582,942,905 | Chr9: 40,769,993 – 43,560,918 |

**Table S6.**
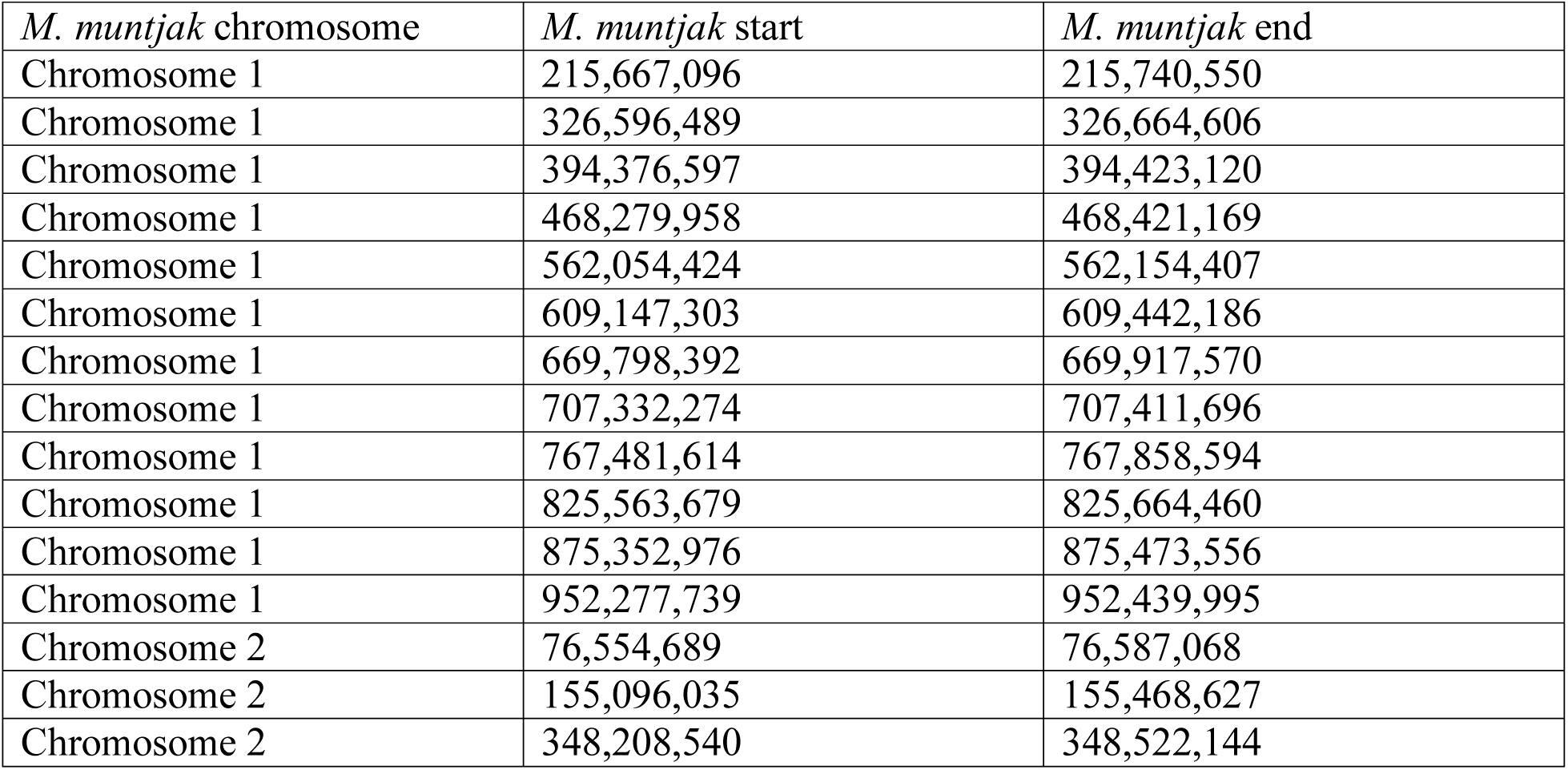
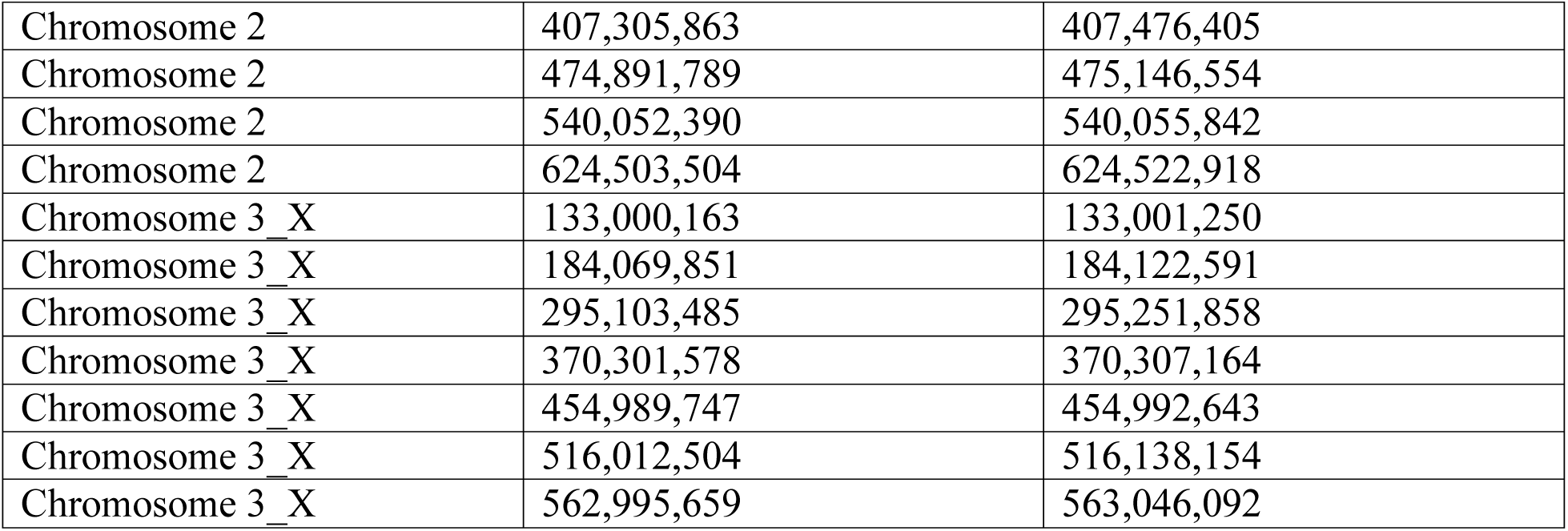
Locations of twenty-six unique fusion events in the *M. muntjak* genome assembly derived from one-to-one orthologs between *M. muntjak* and *M. reevesi* and then refined using runs of collinearity from *B. taurus* and *M. reevesi* against *M. muntjak*.

| *M. muntjak* chromosome | *M. muntjak* start | *M. muntjak* end |
| --- | --- | --- |
| Chromosome 1 | 215,667,096 | 215,740,550 |
| Chromosome 1 | 326,596,489 | 326,664,606 |
| Chromosome 1 | 394,376,597 | 394,423,120 |
| Chromosome 1 | 468,279,958 | 468,421,169 |
| Chromosome 1 | 562,054,424 | 562,154,407 |
| Chromosome 1 | 609,147,303 | 609,442,186 |
| Chromosome 1 | 669,798,392 | 669,917,570 |
| Chromosome 1 | 707,332,274 | 707,411,696 |
| Chromosome 1 | 767,481,614 | 767,858,594 |
| Chromosome 1 | 825,563,679 | 825,664,460 |
| Chromosome 1 | 875,352,976 | 875,473,556 |
| Chromosome 1 | 952,277,739 | 952,439,995 |
| Chromosome 2 | 76,554,689 | 76,587,068 |
| Chromosome 2 | 155,096,035 | 155,468,627 |
| Chromosome 2 | 348,208,540 | 348,522,144 |
| Chromosome 2 | 407,305,863 | 407,476,405 |
| Chromosome 2 | 474,891,789 | 475,146,554 |
| Chromosome 2 | 540,052,390 | 540,055,842 |
| Chromosome 2 | 624,503,504 | 624,522,918 |
| Chromosome 3_X | 133,000,163 | 133,001,250 |
| Chromosome 3_X | 184,069,851 | 184,122,591 |
| Chromosome 3_X | 295,103,485 | 295,251,858 |
| Chromosome 3_X | 370,301,578 | 370,307,164 |
| Chromosome 3_X | 454,989,747 | 454,992,643 |
| Chromosome 3_X | 516,012,504 | 516,138,154 |
| Chromosome 3_X | 562,995,659 | 563,046,092 |

**Table S7.** Locations of six unique fusion events in the *M. reevesi* genome assembly derived from one-to-one orthologs between *M. muntjak* and *M. reevesi* and then refined using runs of collinearity from *B. taurus* and *M. muntjak* against *M. reevesi*.

| *M. reevesi* chromosome | *M. reevesi* start | *M. reevesi* end |
| --- | --- | --- |
| Chromosome 1 | 216,558,007 | 216,594,231 |
| Chromosome 2 | 78,441,940 | 78,562,328 |
| Chromosome 3 | 154,290,838 | 154,298,872 |
| Chromosome 3 | 191,988,183 | 192,099,419 |
| Chromosome 4 | 111,704,918 | 111,709,617 |
| Chromosome 5 | 47,104,391 | 47,224,935 |

